# Transcript accumulation rates in the early *Caenorhabditis elegans* embryo

**DOI:** 10.1101/2021.10.06.463414

**Authors:** Priya Sivaramakrishnan, Cameron Watkins, John Isaac Murray

## Abstract

Dynamic changes in transcription are widespread in developing embryos, where cell cycles are rapid and cell fate decisions need to be made quickly, often before the next cell division. Some fate decisions in the early *Caenorhabditis elegans* embryo overcome these constraints through the rapid production of high absolute levels of transcription factor mRNAs. Single cell accumulation rates for a small subset of developmental genes are known, but genome-scale measurements are lacking. Furthermore, how different aspects of transcription kinetics are fine-tuned for different genes to achieve the appropriate RNA levels is still being worked out. We describe a novel strategy to analyze single cell RNA sequencing data from the early *C. elegans* embryo. We estimate the absolute accumulation rates of zygotic genes up to the 16-cell stage and calibrate predicted rates with single molecule transcript imaging. We show that rapid transcript accumulation is common across different cell types and lineages and rates are the highest soon after zygotic transcription begins. High-rate transcription is a characteristic of genes encoding transcription factors with functions in cell fate specification. These genes share common genomic features and are more likely to have undergone recent duplication. We identify core promoter motifs that might drive high absolute RNA accumulation rates. We measured the contributions of core promoter elements to accumulation rate for one rapidly accumulating gene, *ceh-51*, which is required for mesoderm development. We find that mutating individual motifs modestly decreases the accumulation rate of *ceh-51* mRNA, suggesting multifactorial control of transcript accumulation rates. These results are a step towards estimating absolute transcription kinetics during embryonic fate specification and understanding how transcript dosage drives developmental decisions.

## INTRODUCTION

The embryonic transcriptome is highly dynamic and zygotic transcription in the right cells, at the right time and at the right levels is important for robust cell fate decisions during early development. Absolute transcript accumulation rates (rate of change of RNA copy number) on a global scale are rarely explicitly measured during development. Quantitative imaging of transcriptional activity during development, most extensively in *Drosophila* embryos, has provided detailed measurements of the transcript production rates of early patterning genes (*1-3*). One such study found that the accumulation rates of the gap genes *hunchback, krüppel, knirps* and *giant* to be fairly similar (33 transcripts/minute per nucleus) during the 15-minute nuclear cell cycle 13 (nc13) (*4*). These rates are high, with levels approaching the theoretical steric limit based on the RNA polymerase II (PolII) footprint which would allow for one PolII every 70-80 basepairs. These measurements have been borne out by live imaging of mRNAs, which has provided significant details on the underlying dynamics of rapid accumulation - including the rates of transcription elongation and the response of *cis*-regulatory elements to input maternal gradients (*1, 3, 5*). However, live transcript imaging often relies on transgenes and not the native gene context. Further, most fly studies focus on the early embryo which is syncytial (nuclei sharing common cytoplasm), and it is less clear whether similarly high accumulation rates are common in cellularized embryos and if high rates extend beyond the well-studied patterning genes. Genome-wide bulk RNA sequencing measurements of transcript accumulation on a broad time scale (multiple rounds of cell division) in the *Xenopus* embryo similarly showed that most genes, including transcription factors, had uniform accumulation rates of ∼10^5^ transcripts/hour per whole embryo (*6*). Averaging this accumulation over multiple contributing cells may correspond to a much lower per cell rate, leaving open the question of how rates are controlled in individual cells; and the impact of rate dysregulation on developmental fate decisions remains underexplored.

The *Caenorhabditis elegans* embryo provides an ideal developmental system to answer these questions. It develops with an invariant lineage where division patterns are highly reproducible across embryos, allowing equivalent cells to be compared between individuals (*7, 8*). This robustness results in part from a high level of precision in expression levels of cell fate specification genes, which has been observed for intestinal transcription factors (*9, 10*). The intestinal specification GATA factors *end-3* and *end-1* reproducibly accumulate to very high maximum transcript levels (>300 mRNA molecules per cell) in short (15-20 minute) cell cycles, suggesting that these genes are transcribed at very high rates (*9, 10*). Proper gut formation requires these transcripts to surpass a threshold level, emphasizing the importance of transcription rate in developmental progression (*9, 11*).

Single cell, single molecule transcript imaging methods have been indispensable for visualizing and measuring transcript changes and quantifying the absolute number of mRNA molecules during development (*12, 13*). Single molecule RNA fluorescence *in situ* hybridization (smFISH) in fixed embryos has highlighted the importance of precise transcript levels for developmental robustness (*4, 9, 14*). However, smFISH is not easily scalable for genome-wide interrogation of transcription rates. Studies in cell culture have shown good correlations between reads from different single cell RNA sequencing (scRNA-seq) platforms and smFISH transcript counts, suggesting that scRNA-seq data may allow for genome-wide estimates of absolute transcript levels (*15-17*).

To determine if high transcript accumulation rates are common across early *C. elegans* zygotic genes, and to characterize rates genome-wide, we asked whether read counts in embryonic scRNA-seq data are predictive of absolute transcript levels. Comparing rates inferred from scRNA-seq with smFISH as a gold standard, showed the utility of scRNA-seq in identifying high-rate genes. We found that genome-wide distributions of transcript accumulation rates vary substantially across cell types and lineages, and identified gene features associated with rapid accumulation, including short primary transcript length, and lower intron count. Promoters of high-rate genes include binding sites for lineage-specific transcription factors and are enriched for the Initiator (Inr) motif. Finally, we show that each of these core promoter motifs contributes incrementally to rapid transcription of an exemplar high-rate gene, *ceh-51*. Our results suggest that rates are regulated at multiple levels, consistent with the robustness of *C. elegans* embryogenesis.

## RESULTS

### Analysis of scRNA-seq data reveals differences in transcript accumulation rates in the early *C. elegans* embryo

We predicted genes with high transcript accumulation rates by analyzing an existing scRNA-seq dataset that includes cells up to the 16-cell stage of the *C. elegans* embryo (*18*) (Fig. 1A, see Methods). In *C. elegans*, a first wave of zygotic genome activation occurs at 4-cell stage, leading to the establishment of the main founder lineages by the 16-cell stage. We estimated total RNA counts in each cell by correcting the transcripts per million (TPM) for cell volume, based on previous work showing that mRNA content scales with cell size (*19, 20*), including in *C. elegans* embryos (*18*). The total mass of RNA remains relatively unchanged in the early stages. Additionally, comparing the TPM and smFISH data for the previously studied endodermal genes led us to use an estimate of 8,000,000 mRNA molecules per embryo (*10, 18*), which we also used to calculate the final absolute RNA counts. We used two metrics to identify rapidly accumulating zygotic transcripts. First, we calculated the absolute change for each gene by subtracting the number of transcripts in each cell from those predicted to be contributed by its mother (Fig. 1A, Methods). This provides an estimate of the number of new transcripts produced during each cell cycle. Second, we used the fold-change from parent to daughter to distinguish newly transcribed genes from those already present at high levels in the parent. Finally, we removed maternally expressed genes (TPM >12.5 in P0) to limit our analysis to zygotically transcribed genes.

**Figure 1.**
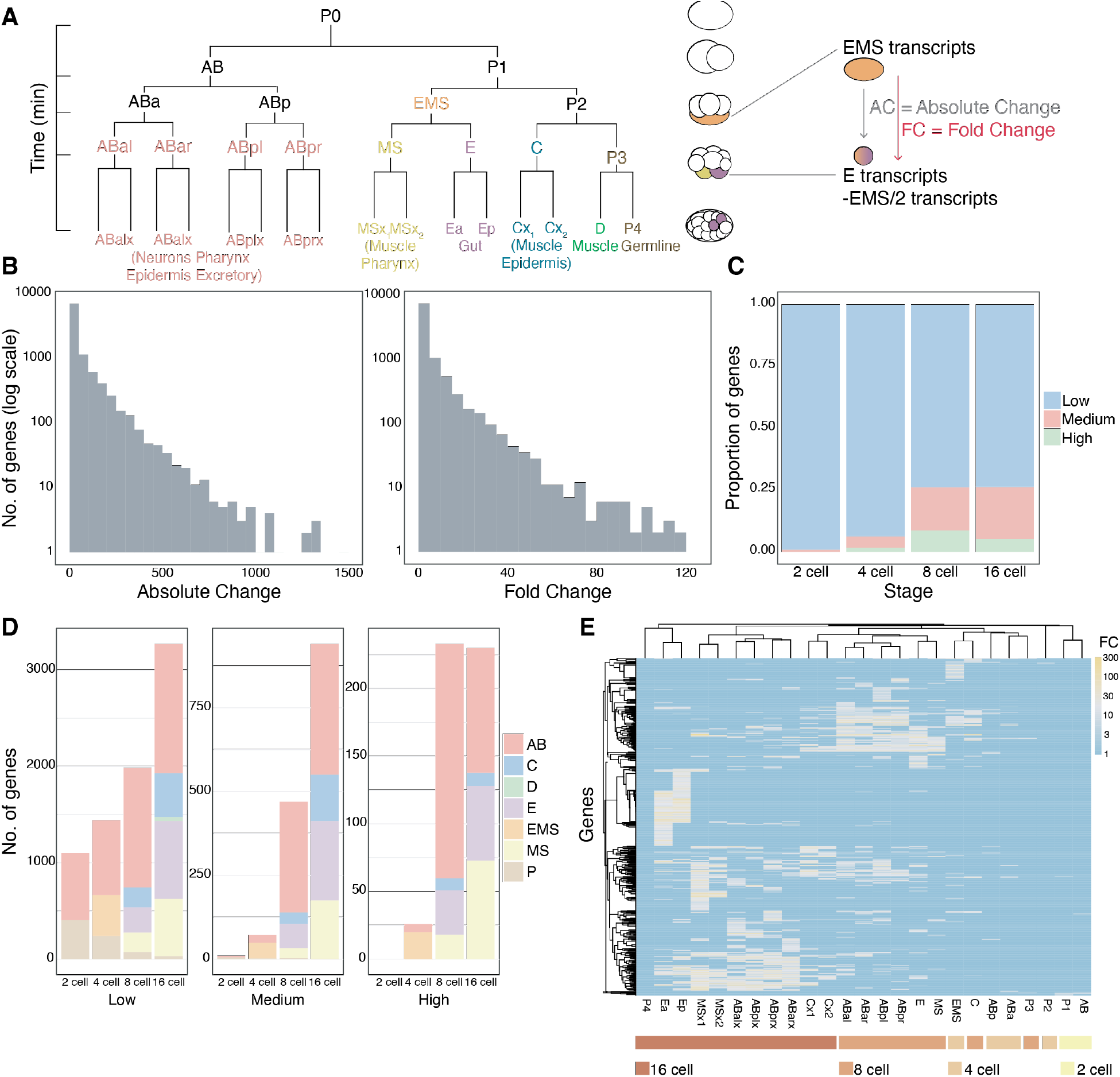
Estimating absolute transcription rates from scRNA-seq data. **A**. Lineage tree of the *C. elegans* embryo up to the 16-cell stage showing all the founder cells (AB, EMS, MS, E, C, D and P) and <u>simplified</u> list of the tissue types that they produce. Accumulation rates were determined by change in the number of transcripts in daughters compared to the parent cell (see Methods). **B**. Histogram distribution of absolute change (AC) and fold change (FC) for all genes and cells. **C**. Fraction of the transcriptome at each embryonic stage based on rate category - high, medium and low. **D**. Number of genes in each rate category by embryo stage and main founder lineages. **E**. Heat map of all high-rate genes (high and medium) using average method of clustering with Pearson distances. Colored bars show embryo stage.

By combining both absolute and fold change metrics, we identified newly transcribed genes with high rates of transcript accumulation. In order to identify common features associated with different rates of accumulation and study rate regulation, we broadly categorized genes based on their absolute and fold change (Fig. 1B). We classified genes based on a loose binning strategy as “high-rate” - having at least 200 new transcripts and FC>10, “medium-rate” - at least 50 new transcripts and FC>5 and “low-rate” - <50 new transcripts and FC<5 (Fig. 1B, Supplementary Fig. 1B). The majority of zygotically expressed genes showed low accumulation rate across all early embryonic stages tested, but a substantial fraction accumulates at high (17.5% at the 8-cell stage and 5.2% at the 16-cell stage) or medium (8.7% at the 8-cell stage and 21.2% at the 16-cell stage) (Fig. 1C, Supplementary Fig. 1D). We confirmed that at these thresholds, the number of genes predicted to have high or medium rates in germline (P) cells was low, consistent with the expected transcriptional quiescence of these cells (Fig. 1D, Supplementary Fig. 1A). In total, we predicted 205 high-rate and 524 medium-rate genes. Since zygotic genome activation in *C. elegans* begins at the 4-cell stage (*21, 22*) most high- and medium-rate genes are expressed at the 8- and 16-cell stages (Fig. 1C, D). At these stages, high-rate transcription is detectable in most somatic cells except the D blastomere, which was collected soon after its division from the germline in this dataset, giving the D cell minimal time to activate transcription (Fig. 1D). Overall, the AB and E lineages have the largest number of genes with high and medium rates, followed by the MS and C lineages (Fig. 1D, Supplementary Fig. 1C,E). As expected, cells hierarchically clustered based on their absolute change of high- and medium-rate genes grouped by embryo stage (Fig. 1E).

### Validation of accumulation rates

Since scRNA-seq measurements can be noisy (*15*), we tested the validity of our rate predictions by an orthogonal method, smFISH. In this method, we targeted multiple oligonucleotide probes to the exons of target mRNAs, allowing single transcripts to be visualized as diffraction limited spots (Fig. 2A) (*12*). The spots can be counted to determine the absolute number of mature transcripts in each embryo. We determined the stage of each embryo by counting DAPI-labeled nuclei (Fig. 2A). We found that all (7/7) candidate high- and medium-rate genes we tested showed corresponding high transcript accumulation by smFISH, (within 2-fold difference in rate) (Fig. 2C-I). The rates estimated by the two methods show a modest positive correlation (r = 0.63, Fig. 2B). One major contribution to the difference likely comes from staging differences, which is more fine-grained in our smFISH measurements. However, this level of correlation is similar to previous comparisons between different scRNA-seq platforms and smFISH counts in melanoma cell lines (*15*). We infer from this analysis that the majority of high-rate genes identified from scRNA-seq are being correctly classified by our approach.

**Figure 2.**
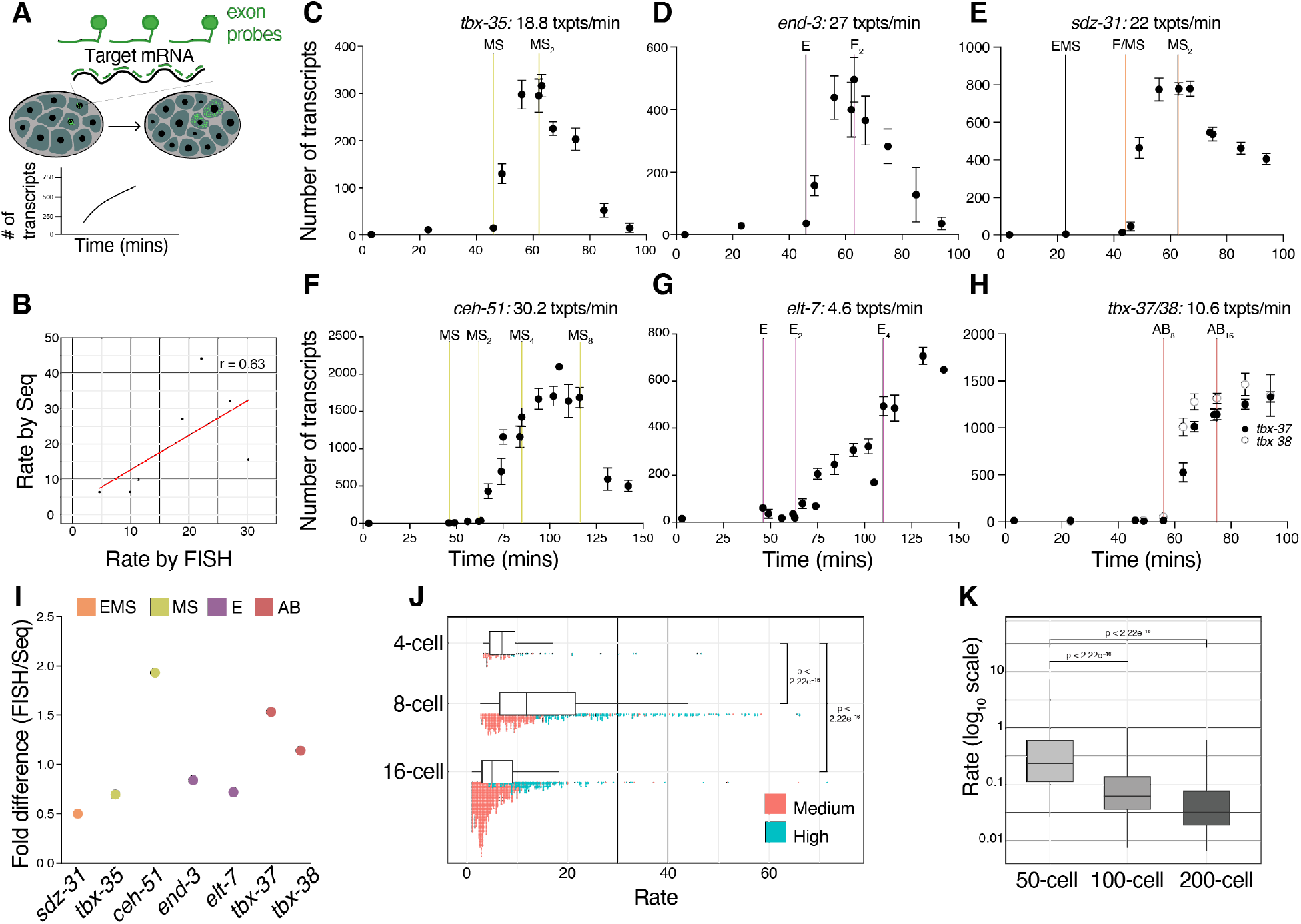
Comparison of inferred rates from sc-RNA seq with smFISH. **A**. Single molecule FISH (smFISH) method used for measuring accumulation rates. Nuclear staining was used to count number of cells to estimate time. **B**. Correlation between estimated rates from scRNA-seq and smFISH (Pearson correlation). **C-H**. smFISH counts (for 7 different genes in different lineages) over developmental time used to calculate transcript accumulation rates between the two cell divisions of maximal expression. **I**. Fold difference in estimated rates by smFISH and scRNA-seq for the 7 genes tested. **J**. Distribution of transcript accumulation rates from scRNA-seq estimates (absolute change/min) for all high- and medium-rate genes across cell types in the AB, EMS, E, MS and C lineages at the 4-, 8- and 16-cell stage. **K**. Accumulation rates at later embryo stages estimated from the Packer et. al data set (*33*). Adjusted *p-*values from Wilcoxon test.

### Maximum transcription rates vary across stages

*C. elegans* embryonic cells change dramatically at each subsequent stage. The embryo undergoes reductive cleavage, with cell and nuclear volumes decreasing by up to ∼2-fold each division (*23*). Meanwhile, cell cycle length increases over time, with each cell dividing slower than its mother (*24, 25*). Finally maternal repressors prevent transcription at the 1-cell and 2-cell stages, and the degradation of these repressors over time facilitates the onset of zygotic transcription at different times in different lineages (*26*). This raises the question of whether maximum transcription rates vary by stage or lineage.

We first examined transcriptional rates of the genes measured by smFISH that are expressed in the MS (mesoderm) and E (intestine) lineages (Fig. 1A). The MS cell expresses *tbx-35*, which encodes a T-box transcription factor required for normal mesodermal development (*27*). Between the birth and the division of the MS cell (16 min), ∼300 transcripts of *tbx-35* accumulate, giving an accumulation rate of 18.8 transcripts/minute (Fig. 2C). *ceh-51*, which is expressed one cell cycle later in the MS daughter cells and is activated by TBX-35 (*28*), accumulates at an even higher rate of 30.2 transcripts/minute (Fig. 2F). *end-3*, which encodes a GATA transcription factor involved in intestinal fate specification is expressed in the E cell but at the same time as *tbx-35* has a similarly high rate of 27 transcripts/minute (Fig. 2D). However, *elt-7*, a main target of *end-3* in the E daughter cells (*29*), has a much lower accumulation rate (4.5 transcripts/min; Fig. 2G). Even though *elt-7* shows a fairly high final transcript level (∼700 total transcripts), this occurs over the E2-E4 division which has a much longer cell cycle length. The transcripts of *sdz-31*, a predicted membrane protein (*30*), accumulates in both E and MS cells at 22 transcripts/min per cell (Fig. 2E). Finally, we also imaged *tbx-37* and *tbx-38*, two paralogous genes expressed early in the ABa lineage and play important fate specification roles in ABa descendants (*31, 32*). *tbx-37/38* accumulate at ∼11 transcripts/min in ABa daughters (Fig. 2H). Our smFISH analysis suggests that RNA accumulation rates can vary from gene to gene and that the AB, E and MS lineages are capable of high-rate transcription. However, higher rates are seen earlier (8-cell stage) when compared with the 16-cell stage.

To ask whether there are genome-scale differences in transcript accumulation rate between stages and lineages, we calculated the accumulation rate of all high- and medium-rate genes detected in the scRNA-seq data. We found that while high-rate transcription occurs at the 4-cell, 8-cell and 16-cell stages, concordant with our smFISH imaging, the distribution of rates was highest at the 8-cell stage (Fig. 2J). This seems broadly true across most lineages; the median accumulation rate of medium- and high-rate genes was higher for cells at the 8-cell stage compared to their daughters at the 16-cell stage (Supplementary Fig. 2A). Overall median rates at the 8-cell stage are ∼3-fold higher compared to the 4-cell (Supplementary Fig. 2B). To contrast this with later development, we estimated the absolute rate change in later cells from the scRNA-seq data set generated by Packer et al. (*33*). The estimated rates at the 8-cell stage are 49-fold higher than the rates for cells born around the 50-cell stage, and rates continue to decline at the 100- and 200-cell stage (Fig. 2K). Within these later stages, rates are fairly consistent across lineages (Supplementary Fig. 2C). We conclude that the extent of high-rate transcription varies between embryo stages, with an apparent peak at the 8-cell stage, very quickly after the onset of zygotic transcription.

### High-rate genes include dosage-sensitive transcription factors (TFs)

To understand the types of genes that are rapidly transcribed and determine if they are involved with developmental fate decisions, we performed Gene Ontology analysis on high- and medium-(high + medium) and low-rate genes to identify categories of biological process and molecular function associated with the two rate groups. High- and medium-rate genes were enriched for annotations associated with transcription factors (DNA binding activity) and development (cell fate specification and gastrulation) (Fig. 3A). By contrast low-rate genes are enriched for terminal cell type-specific annotations and housekeeping functions (Fig. 3A).

**Figure 3.**
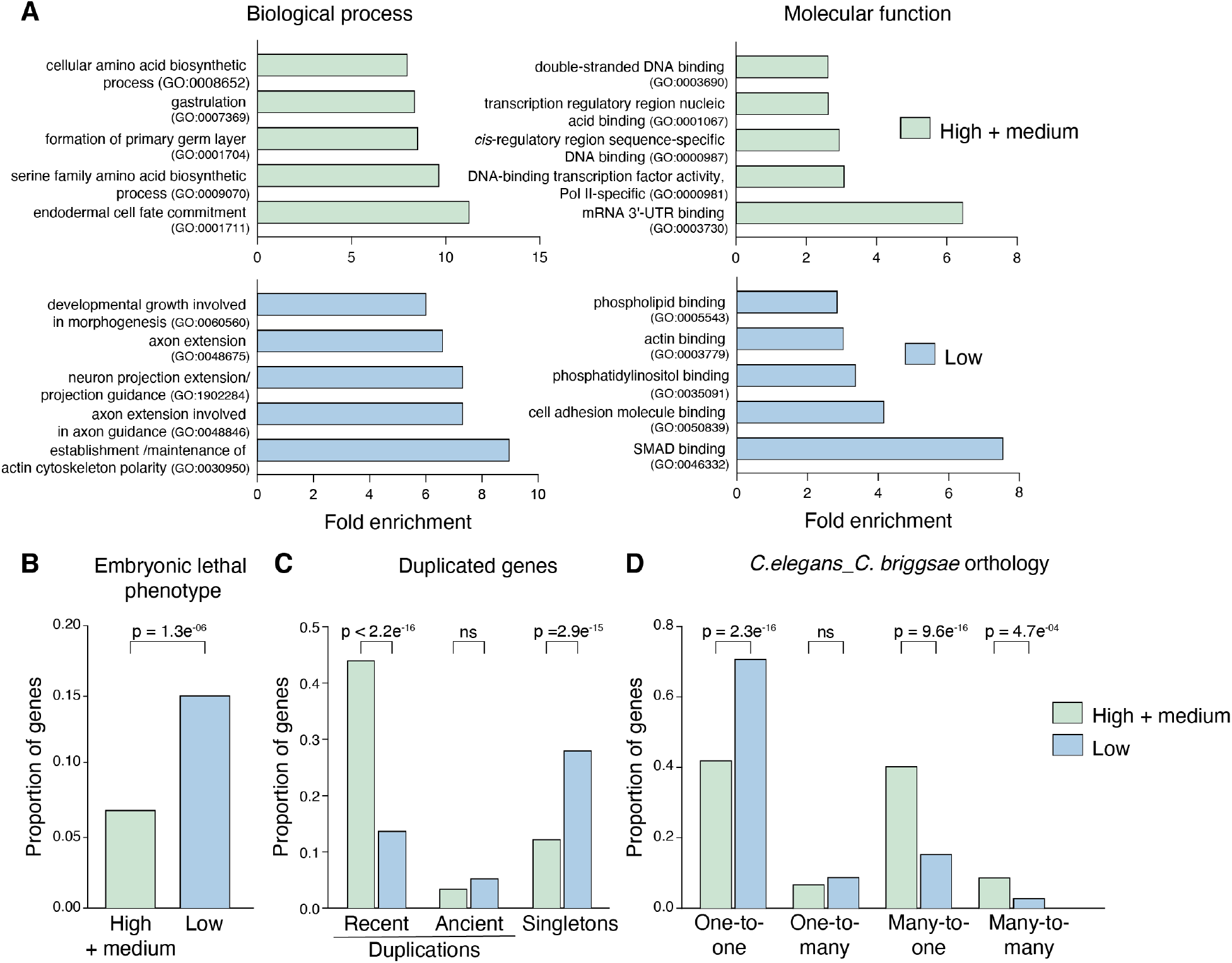
Function and evolutionary conservation of high vs. low-rate genes. **A**. Gene ontology categories that show enrichment within high and medium (combined) and low-rate genes, from PANTHER analysis. Top 5 GO terms shown for each, representative terms were used for similar categories. **B**. Proportion of all high and low-rate genes with embryonic lethal phenotypes. **C**. Proportion of all high and low-rate genes that have duplications or are singletons (using data from Ma et al.(*48*)). **D**. Orthogroups identified between *C. elegans* and *C. briggsae* in all high vs. low-rate genes. All reported *p-* values from Fisher’s exact test.

We asked whether rapidly transcribed genes were more likely to be essential by checking if there were associated with known embryonic lethal mutant or RNAi phenotypes as curated by WormBase (*34*). Surprisingly, the proportion of low-rate genes with embryonic lethal phenotypes was significantly greater than the high- and medium-rate genes (Fig. 3B). Since several of the high-rate genes we examined by smFISH (*end-3, elt-7, tbx-37*) have partially redundant paralogs (*35*), we wondered if widespread redundancy due to gene duplications could mask the phenotypes of high-rate genes. Consistent with this, high-rate genes are significantly more likely to have been recently duplicated (Fig. 3C, 3-fold enriched compared with low-rate) and have a predicted paralog (Supplementary Fig. 3A), while more low-rate genes occur as singletons (Fig. 3C).

Since high-rate genes were more likely to have recent duplicates in *C. elegans*, we asked whether they were also more likely to have multiple orthologues in *C. briggsae (36*). Although they diverged ∼30 million years ago, both species are morphologically and behaviorally similar (*37*). We found that low-rate genes were 1.7-fold more likely to have one-to-one orthologs in *C. briggsae* compared to high- and medium-rate genes (Fig. 3D). However, a greater proportion of high- and medium-rate genes have many-to-one (2.5-fold over low-rate) or many-to-many orthologs (3-fold over low rate) in *C. briggsae* (Fig. 3D).

We hypothesized that since low-rate genes are associated with embryonic lethality, they may be more likely to be evolutionarily conserved over longer distances. Using the Ortholist2 (*38*) database of worm-human orthologs, we found that a significantly greater proportion of low-rate genes had a one-to-one human ortholog compared to medium- or high-rate genes (Supplementary Fig. 3B). We conclude that high-rate transcription is associated with increased rates of gene copy number evolution such that two or more of the duplicated paralogous genes need to be mutated to see a phenotype.

### High-rate genes share gene-specific structural features and form genomic clusters

Previous studies in other organisms have shown that zygotic genes tend to be shorter and have shorter introns compared to maternally expressed genes (*39*). We tested whether this was true for highly transcribed zygotic genes in *C. elegans* embryos and found that that high- and medium-rate genes have shorter primary transcript length than low-rate genes (Fig. 4A). The presence of introns has been linked to higher transcription efficiencies (*40, 41*). We found that higher rate is also associated with shorter introns (Fig. 4B). We calculated the number of introns for each rate category (adjusted for gene length) and found a decreasing trend between rate and intron number (Fig. 4D). Our findings are more consistent with the intron delay hypothesis during development, which posits that longer introns preclude high levels of expression in early development when cell cycles are short (*42, 43*). We also found that high- and medium-rate genes tend to have fewer annotated isoforms (Fig. 4E).

**Figure 4.**
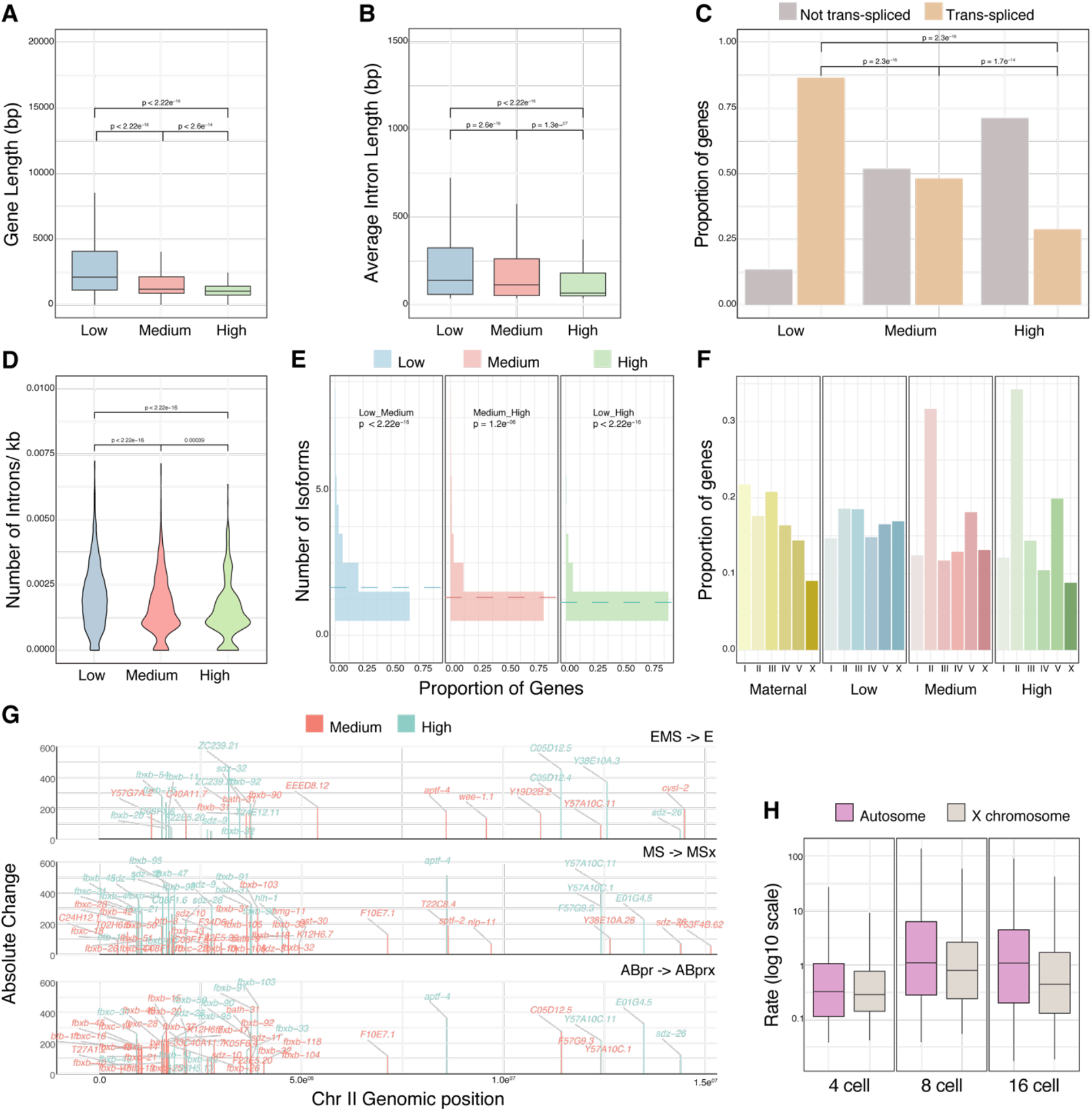
Genetic architecture and features associated with high transcript accumulation. **A**,**B**. Gene and intron lengths across different rate categories. **C**. Proportion of genes that are trans-spliced or not in each rate group using data from Saito et al. (*45*). **D**. Length normalized intron number in each rate category. **E**. Distribution of isoform number, based on proportion of genes in each rate category. Dotted line shows mean. All reported significance from Wilcoxon test with adjusted *p*-values. **F**. Number of genes (only protein coding included) as a proportion of the genes in each category found on the different chromosomes. **G**. Karyogram of genes with FC >5 on chromosome II showing absolute change for each gene across genomic position in the indicated cells compared to their parent. Specificity score indicates how specific the expression is to the particular cell (purple is specific to that cell, red is broadly expressed in all cells). **H**. Accumulation rates by stage and type of chromosome (X vs. autosome).

Around 70% of *C. elegans* genes undergo *trans*-splicing, where the 5’ end of RNAs are replaced with a splice leader that facilitates translation (*44*). We mined existing *trans*-splicing data (*45*) and found that high and medium-rate genes are significantly less likely to be *trans*-spliced (Fig. 4C, ∼3-fold difference between high and low-rate genes).

Position effects, where chromosomal position influences gene expression, are widespread in many species (*46*). To begin understanding the impact of chromosomal position on transcript accumulation rates, we analyzed the distribution of genes in each rate category across chromosomes. We found an over-representation of high and medium-rate genes on chromosome II (Fig. 4F). Examining the position of high-rate genes within each chromosome shows large clusters of rapidly transcribed genes on the very left arm of chromosome II (Fig. 4G, example cells E, MSx1 and ABprx). Most of these clustered are either in the F-box family or the *bath* (BTB and MATH domain containing) family, both of which are substrate-binding adapters for ubiquitin-mediated proteolysis and are rapidly evolving in *Caenorhabditis (47*). More clusters of genes are seen on chromosome II compared to other chromosomes in several cell types, perhaps reflecting the large cluster of F-box genes that is enriched on this chromosome (*48*) (Supplementary Fig. 4). Finally, we see that accumulation rates are higher for genes on autosomes compared with the X chromosome, consistent with reduced expression from the sex chromosome in the germline and early embryo (Fig. 4H).

These results indicate that high-rate transcript accumulation is correlated with specific gene architectures and genomic location. Whether chromosomal clustering, the absence of *trans*-splicing, shorter genes and fewer isoforms mechanistically facilitate high-rate transcription or are correlated for other reasons should be explored further in the future.

### The Inr motif is enriched in promoters of high-rate genes

We next asked what *cis*-regulatory elements play a role in controlling accumulation rates. The minimal genome of *C. elegans* necessitates transcriptional regulation through *cis*-regulatory elements close to the transcription start site (TSS) (*49*). In the developing embryonic gut, most fate specification factors are largely controlled by the promoter-proximal regions rather than distal enhancers (*50*). To find promoter motifs that might control rapid transcription across different cell types, we first combined three different datasets (*45, 51, 52*) to identify the most likely TSS (see Methods). We then asked what sequence motifs were enriched in high- and medium-rate gene promoters (operationally defined as the 500 base pairs upstream of the TSS). We tested both motifs identified by the *de novo* motif finding program MEME, which included the well-studied core promoter motifs Inr and SP1, also the TATA box (Fig. 5A). We also examined the binding sites for known early lineage-specific TFs (END, MED, POP-1, SKN-1 and TBX) (Fig. 5B).

**Figure 5.**
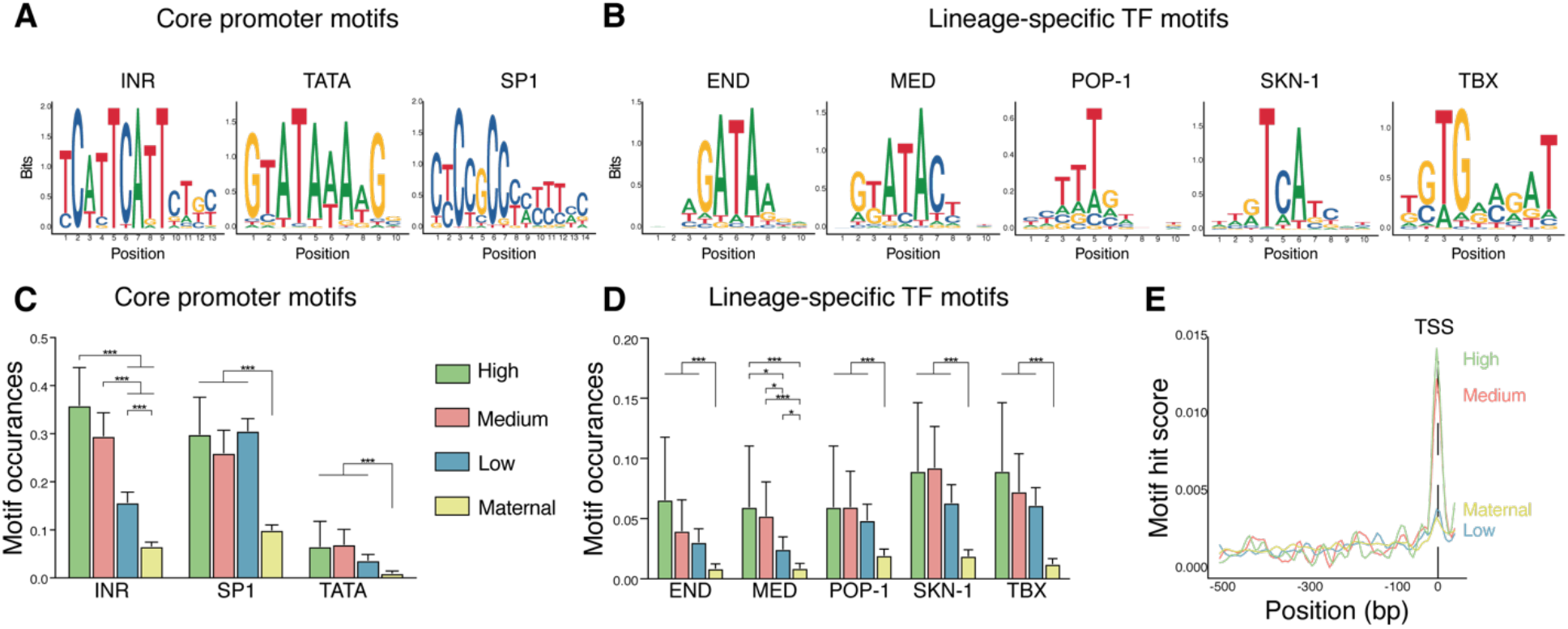
Promoter motifs associated with rapid transcription. **A**,**B**. Position scores of core promoter (A) and lineage-specific motifs (B). Inr and SP1 motifs from the MEME suite, TATA motif from Chen et al. (*51*) and lineage-specific TF motifs from CIS-BP (http://cisbp.ccbr.utoronto.ca/). **C**,**D**. Motif occurrences for the core promoter elements in A (C) or lineage-specific TFs (D) across different rate categories and maternal genes, 95% confidence intervals as error bars, *p-*values from chi-squared test, FDR adjusted *p-*values. **E**. Inr motif distribution across the promoter (500bp upstream) in genes from the indicated rate categories showing overlap of Inr with the TSS.

We observed that binding sites for lineage-specific regulators were enriched in high-rate promoters active in the expected cells. For example, the SKN-1 binding site was enriched in high-rate genes expressed in the EMS cell (Supplementary Fig. 5A) and the MED and END motifs were more likely to be found in promoters active in the E cell or its daughters (Supplementary Fig. 5B). Also as expected, binding sites for these regulators are mostly found in early embryonic genes across all three rate categories in contrast with maternal genes (Fig. 5D).

Among the core promoter motifs, the initiator element (Inr) stood out as strongly associated with rate. Transcription of many *C. elegans* genes has been previously shown to initiate at the ‘A’ of the tcAttc core Inr motif (*51*). Inr motifs had significantly greater occurrence in high-rate genes compared with medium-rate genes, and in both categories compared to low-rate and maternal genes (Fig. 5C). Further, the Inr motif was strongly enriched at the TSS of high and medium-rate genes, but this enrichment was lower for low-rate or maternal genes (Fig. 5E). The TATA box and binding sites for a canonical general transcription factor SP1, are highly but uniformly enriched in low, medium and high-rate genes relative to maternal genes (Fig. 5C), suggesting that distinct core promoter motifs may be used for zygotic *vs*. maternal gene expression.

### Multiple promoter motifs contribute to accumulation rates

We used the high-rate gene *ceh-51* as a model to test the importance of TSS-proximal motifs for transcript accumulation rates. The T-box transcription factor TBX-35 is known to regulate *ceh-51* and four binding sites for TBX-35 were previously identified in the upstream region (promoter) of *ceh-51* (*28*) (Fig. 6A). Deletion of 2 of the TBX-35 binding sites on a CEH-51::GFP extrachromosomal array reduced protein levels, but removal of all 4 sites completely abrogated expression at the MS8 cell stage (*28*). We used CRISPR to mutate TBX-35 binding sites in the endogenous *ceh-51* promoter. We obtained a mutant carrying a 14bp deletion that disrupts the core of the third TBX-35 site; and used smFISH to determine if the accumulation rate of *ceh-51* was affected by this deletion. We measured counts of *elt-7*, which is expressed at the same time as *ceh-51* but in the E cells as a control in each embryo. Loss of this TBX-35 sites resulted in a 1.6-fold reduction in *ceh-51* accumulation from the 14-cell to the 26-cell stage, the stage when *ceh-51* has its maximal accumulation rate (Fig. 6C). Another mutant that disrupts two TBX-35 sites (sites 3 and 4) along with an intervening SP1 site (Supplementary Fig. 6A), had 25% reduction in *ceh-51* rate (Fig. 6B, C). The *C. elegans* homolog of SP1, SPTF-3, is a general transcription factor that has been previously shown to be required for proper expression of gut specification genes and the regulation of gastrulation (*53*). Our observation that SP1 sites are enriched in all zygotic rate classes relative to maternal genes (Fig. 5C), suggests that SP1 may be especially important during embryonic development. Consistent with this, knockdown of maternal *sptf-3* by RNAi leads to partially penetrant embryonic and larval arrest (Supplementary Fig. 6B). However, *sptf-3* RNAi did not significantly change the viability of the TBX-35 site 3,4 SP1 site 1 mutant (Supplementary Fig. 6B).

**Figure 6.**
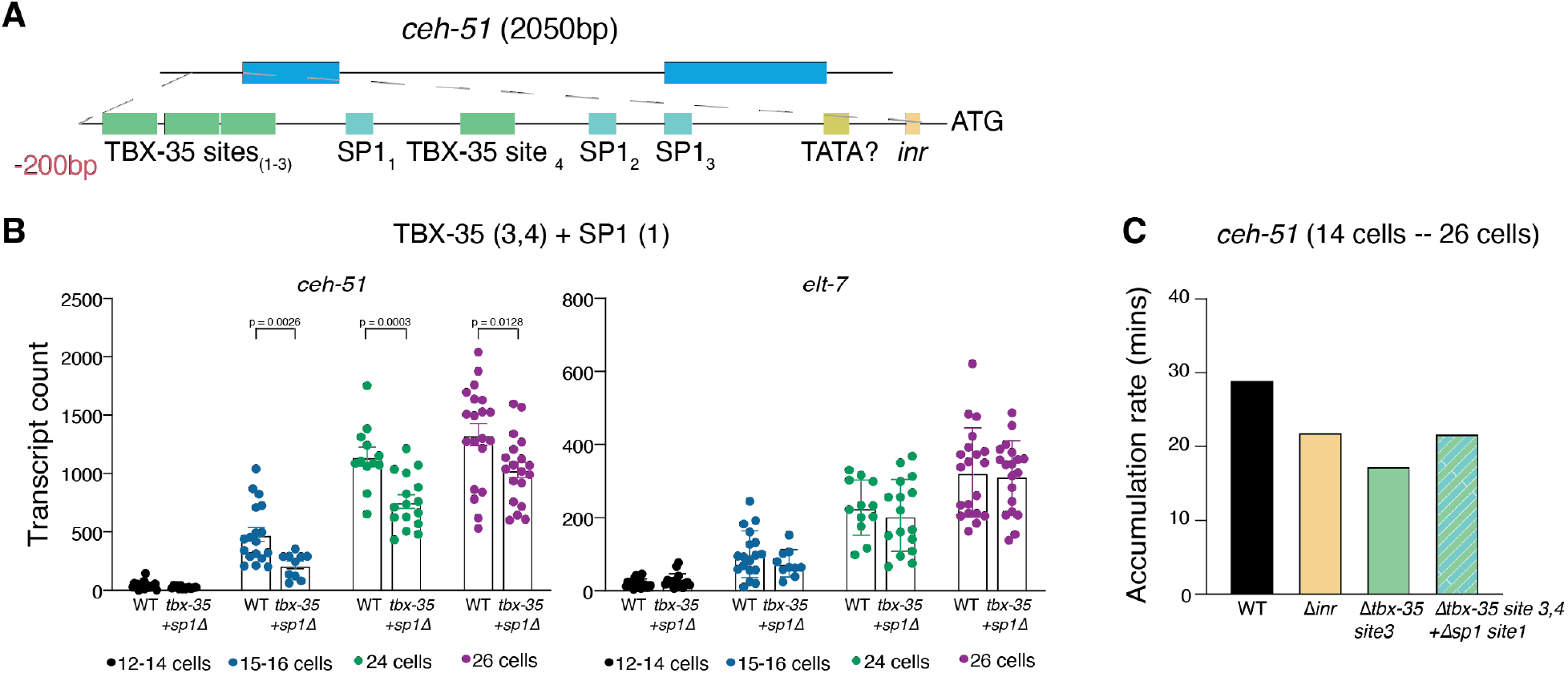
Contribution of promoter motifs to regulation of *ceh-51* transcript accumulation rate. **A**. 200bp upstream region (from translation start ATG) of *ceh-51*. **B**. smFISH exon counts of *ceh-51* and *elt-7* at the indicated stages in the promoter mutant where TBX-35 sites 3 and 4 and SP1 site 1 are deleted, n = at least 3 biological replicates, *p-*values from Mann-Whitney test. **C**. Accumulation rate of *ceh-51* (from the 14 cell stage - 26 cell stage) in the indicated mutants, data from at least 3 biological replicates.

Based on the enrichment and close overlap of the Inr motif with the TSS of high-rate genes, we expected that the Inr may not only be required for rate control but also for overall expression of high-rate genes. To test this, we deleted the core motif of the Inr element in the *ceh-51* promoter and surprisingly found only a 24% reduction in the *ceh-51* accumulation rate (Fig. 6C). Embryonic survival is unaffected in *ceh-51* promoter mutants in which the Inr or a single TBX-35 binding site is disrupted (Supplementary Fig. 6C). Thus, it appears that high rates of transcript accumulation in the early embryo is under partially redundant control, such that not a single motif but a combination of *cis*-regulatory elements together specify the rapid accumulation of *ceh-51*.

## DISCUSSION

### scRNA-seq as a tool to predict absolute transcript levels

Our comparisons with smFISH counts suggest that the estimation of absolute transcript levels can be inferred from scRNA-seq data with reasonable accuracy. Similar correlations, with disagreements typically in the +/- 2-fold range, were seen in previous comparisons of smFISH and scRNA-seq in human cell lines, suggesting they may be generalizable across species and cell types (*15*). In fact, the actual agreement may be higher as we can only infer the correspondence between the stages analyzed by scRNA-seq and smFISH within a few minutes using nuclear counts. Thus, the correlation between imaging and sequencing counts during development could potentially be improved by more precise staging and collection of cells for sequencing. Recently developed high-throughput FISH methods such as seqFISH+ or MERFISH, which can image up to 10,000 genes, report strong correlations between single molecule transcript counts and RNA seq (*16,54*). However, imaging methods still require the design of expensive probe sets. Thus, comparing smFISH counts and scRNA-seq reads or unique molecular identifiers (UMIs) in multiple model systems should help develop principles for inferring absolute transcript levels from sequencing data.

### Importance of high transcript accumulation rates during embryonic development

Appropriate dosage of regulators is important for cell fate decisions during *C. elegans* embryonic development (*9,11*). Transcription factor (TF) dosage is similarly crucial for embryonic patterning in *Drosophila (13,55*) and in mammalian development and is slowly being recognized as underlying causes of human disease (*56*). We predict that transcript accumulation rates are key to achieving the final dosage of TFs within a critical time window, allowing for robust fate specification. We find that many high-rate genes are more likely to have been recently duplicated suggesting that they have paralogs with partially redundant functions (Fig. 3C, Supplementary Fig. 3A). These results are consistent with the hypothesis that rapid transcription occurs to increase the transcript dose for TFs involved in cell fate decisions. One explanation for the tendency of these TFs to have co-expressed paralogs is that the required functional dosage of transcripts is not easily achievable from a single gene, with combined transcripts from paralogous genes required to reach the appropriate threshold level. The significant enrichment of high-rate genes in the many-to-many orthology group suggests that the need for duplicate genes to achieve rapid transcription might be conserved in *C. briggsae*, which could lead to independent gene duplications (Fig. 3E). We also find that clusters of high-rate genes (for example on chromosome II) belong to the same gene family, which likely also has risen out of duplication (*48*) (Fig. 4F). Genes involved in gut specification, such as the *med-* and *end-* GATA factors, tend to exist as multiple paralogs across evolution in the *Elegans* supergroup and these genes also exhibit synteny (located in proximity on the same chromosome) across species (*50*). Whether the spatial organization of high-rate genes is required for their expression or is instead a result of the tendency of duplicated genes to form local clusters is an important question to be addressed in the future.

Absolute transcript levels are representative of both transcript production and degradation. In cell culture systems, during the response to stimuli such as lipopolysaccharides or hypoxia, changes in transcript synthesis contribute more for overall changes in RNA levels than do changes in degradation (*57,58*). Our measured accumulation rates reflect both new transcription and RNA degradation; thus, the *de novo* transcription rates may be higher than the observed accumulation rates, making this accumulation even more impressive. Many of the regulators we analyze here are transiently expressed, with most transcripts disappearing within one or two cell cycles of the time of maximal expression. Similarly, some low-rate genes may be transcribed at high rates if their degradation acts to keep their transcripts levels from changing dramatically.

### Cell size and transcription rates

An important normalization for scRNA-seq counts to estimate absolute mRNA numbers is to account for cell volume. Transcription rates are also known to scale with cell size across organisms (*20,59*). The *C. elegans* embryo develops by reductive cleavage in a constant-volume eggshell with cells reducing ∼2-fold in size with each developmental cell division. We see the highest maximal transcript accumulation rates in the 8-cell stage, where cells are larger compared to the 16-cell stage, suggesting that transcription rates in the *C. elegans* embryo may also be driven by cell size in this context (Fig. 2J). Future work should focus on identifying mechanisms involved in this scaling and how the balance between reduction in cell size and the ramping up of zygotic transcription is achieved as early development progresses.

### Boundaries on the maximum achievable transcription rates

What are the theoretical limits on transcription rates? Several factors contribute to this, including the available number of PolII and other transcription factor molecules, the rates of initiation and elongation, gene length and pause sequences. For short genes such as *end-3* (∼1.3kb primary transcript), at the commonly assumed elongation rate of 1.5kb/min (*60, 61*), the entire gene length would need to be completely occupied by PolII complexes during maximum expression. smFISH analysis of early zygotic genes in the *Drosophila* embryo have also measured high rates of transcription, potentially nearing the steric limits of PolII, suggesting that rates approaching theoretical maximum could be a common feature of gene expression in rapidly dividing embryos (*5, 62*). Elongation rates have not been directly measured in the *C. elegans* embryo although elongation speeds as high as 6kb/min are seen in mouse embryonic stem cells (*60*). It will be interesting to determine whether initiation and elongation rates are coordinately high for genes with high accumulation rates.

We have characterized transcription accumulation rates in a developing multi-cellular organism applying a novel approach to analyzing scRNA-seq data and find that the regulation of rates is highly complex. We hypothesize that high rates are required to achieve precision in final transcript levels, which drives fate specification. How this precision will translate to protein levels is still unclear. Our study predicts that both RNA and protein levels ultimately play a role in regulating cell fate specification and overall developmental robustness.

## MATERIALS AND METHODS

### Accumulation rates from scRNA-seq data

All analysis was performed using the R statistical programing language. Analysis code can be found at: github.com/p-sivaramakrishnan/C-elegans-rate-analysis. Gene names from the Tintori et al. dataset were transferred to WS260 reference. We note that since the timing of division varies between lineages, the 16-cell stage in the Tintori dataset reflects the 16 cells of the AB lineage or the 24-cell stage of the embryo. For the sake of simplicity, this stage is still referred to as the 16-cell stage.

Median Transcript per million (TPM) counts for each gene in each cell was calculated, cells annotated as “tossed” were not used. Absolute change for each gene in each cell was calculated as (TPM_daughter_ minus TPM_parent_) * volume of the daughter cell * 8,000,000. The 8,000,000 mRNA molecules per embryo is based on previous smFISH absolute counts measured for endodermal genes (*9, 10*) and unchanging total mass of RNA for each early stage and determined from spike-in controls by Tintori et al. (*18*). Thus, the absolute change metric reflects difference in transcript copy number between parent and daughter, adjusting for pre-existing transcripts inherited by each daughter from the parent. Fold change between daughter and parent was calculated with a pseudocount of 10 (to get estimates even when TPM in the parent was 0). Genes with absolute transcript amounts > 100 (TPM 12.5) in the P0 cell were considered to be maternal genes.

### Rate categorization

Distributions of absolute change (AC) and fold change (FC) (Fig.1B) were used to bin rate categories. AC >200, FC > 10 was considered high-rate, AC between 50-200, FC between 5-10 as medium-rate and AC between 1-50 and between FC 1-5 as low-rate. If the AC or FC was high but the other fell into the medium category, it was classified as medium-rate. Similarly, if AC and FC were medium or high, but FC was low (1-5), those genes were classified as low-rate. For gene structure and motif analysis, only high and medium-rate genes that were never low-rate in any cell were included in the analysis.

### Gene Ontology and conservation with *C. briggsae*

GO was performed using PANTHER tools (http://geneontology.org/) using the Overrepresentation Test (Released 20220202) against the *C. elegans* reference list and Fisher’s exact test with FDR correction. OrthoFinder was used to examine conservation between *C. elegans* and *C. briggsae (36*). Longest transcript isoforms were used to identify Orthogroups, which were then categorized as having one-to-one, one-to-many, many-to-one or many-to-many orthologous genes between *C. elegans* and *C. brissgae*.

### Paralogous genes and embryonic lethal phenotypes

Data on gene duplications was obtained from Ma et al (*48*). A list of paralogs (Blast result from a e-value threshold of 10^−15^) that were also syn-expressed was obtained from Tintori et al. (*18*). Wormbase SimpleMine was used to identify allele and RNAi mutant phenotypes (https://wormbase.org//tools/mine/simplemine.cgi).

### Transcription start site (TSS) refinement

We collated TSS data from three different data sets (Chen et al., Saito et al. and Kruesi et al. (*45, 51, 52*), where TSS was predicted by different methods. This allowed us to predict the most likely single TSS for each gene. Promoter regions were then defined as 500bp upstream of this newly consolidated TSS.

### Motif tools

Xstreme (Meme suite, https://meme-suite.org/meme/) analysis using a background Markov order of 2 and Homer tools (background adjusted to *C. elegans* promoters) were used for *de novo* motif prediction. The Find Individual Motif Occurrences (FIMO) tools in the Meme suite was used for motif enrichment using a p-value cut-off of 1E^-4^ and the R package ggseqlogo was used to plot motifs.

### Worm stains and growth

Worms were grown on standard NGM plates on OP50 bacteria. 1mM IPTG and 50μg/ml carbenicillin were added for RNAi plates. For smFISH, worms were grown on large, enriched peptone plates seeded with NA22 bacteria. All growth was at 20°C unless otherwise stated.

### Single molecule RNA FISH (smFISH)

smFISH was performed as described in Nair et al. (*10*). Briefly, adult worms were treated with alkaline bleach to release embryos, which were washed with M9 buffer. Embryos were fixed with 4% formaldehyde (in PBS) and permeabilized by immersing the tube in dry ice and ethanol bath for 1 minute. After thawing and a 20-minute incubation on ice, embryos were washed with PBS and stored in 70% ethanol at 4°C. For hybridization with FISH probes, first the ethanol was washed off the embryos with wash buffer (10% formamide in 2X SSC with 0.1% Triton X-100). FISH probes were used at 2-4nM concentration in 100μl of hybridization buffer (0.1% dextran sulphate, 0.1% formamide in 2X SSC buffer). Hybridization was performed overnight at 37°C. Before imaging, embryos were stained with Hoechst stain for 30 minutes and mounted on a cover slip in 2X SSC. smFISH images were taken with a standard epifluorescence scope.

### smFISH transcript counts

TIFF stack images were processed using Matlab. Embryos were segmented and a Laplacian of Gaussian filter was applied to isolate single transcripts. Transcript counts were thresholded manually to obtain spot counts across all relevant Z stacks. Each channel for different probe sets were separately thresholded. Hoechst stain was used to determine the number of nuclei and cell stage.

### CRISPR deletions

Promoter deletions were performed by CRISPR using short single-stranded guide RNAs (as described in Dokshin et al. (*63*)). Guides and tracrRNA were obtained from Integrated DNA Technologies (IDT). *S. pyogenes* Cas9 was purchased from the QB3 MacroLab, UC Berkley. Co-injection marker *punc*-119::GFP was used to screen for positive injections. GFP positive F1s were singled and deletions were confirmed by sequencing.

## ACKNOWLEDGEMENTS

We are extremely grateful to Arjun Raj and members of the Raj lab for their generosity in letting us use their microscopes, smFISH troubleshooting, advice on image analysis and overall, for being fantastic neighbors. We would also like to thank the members for the Murray Lab (past and present) and the Penn Worm Group for their insightful discussions and comments.

Some strains were provided by the CGC, which is funded by NIH Office of Research Infrastructure Programs (P40 OD010440). We thank WormBase, which is supported by NHGRI grant U24 HG002223, for consolidating valuable information.

## DATA AND CODE AVAILABILITY

Code is available on Github (github.com/p-sivaramakrishnan/C-elegans-rate-analysis). Source data are available in the supplementary materials.

