## Supplementary figures for "Transcript accumulation rates in the early *Caenorhabditis elegans* embryo"

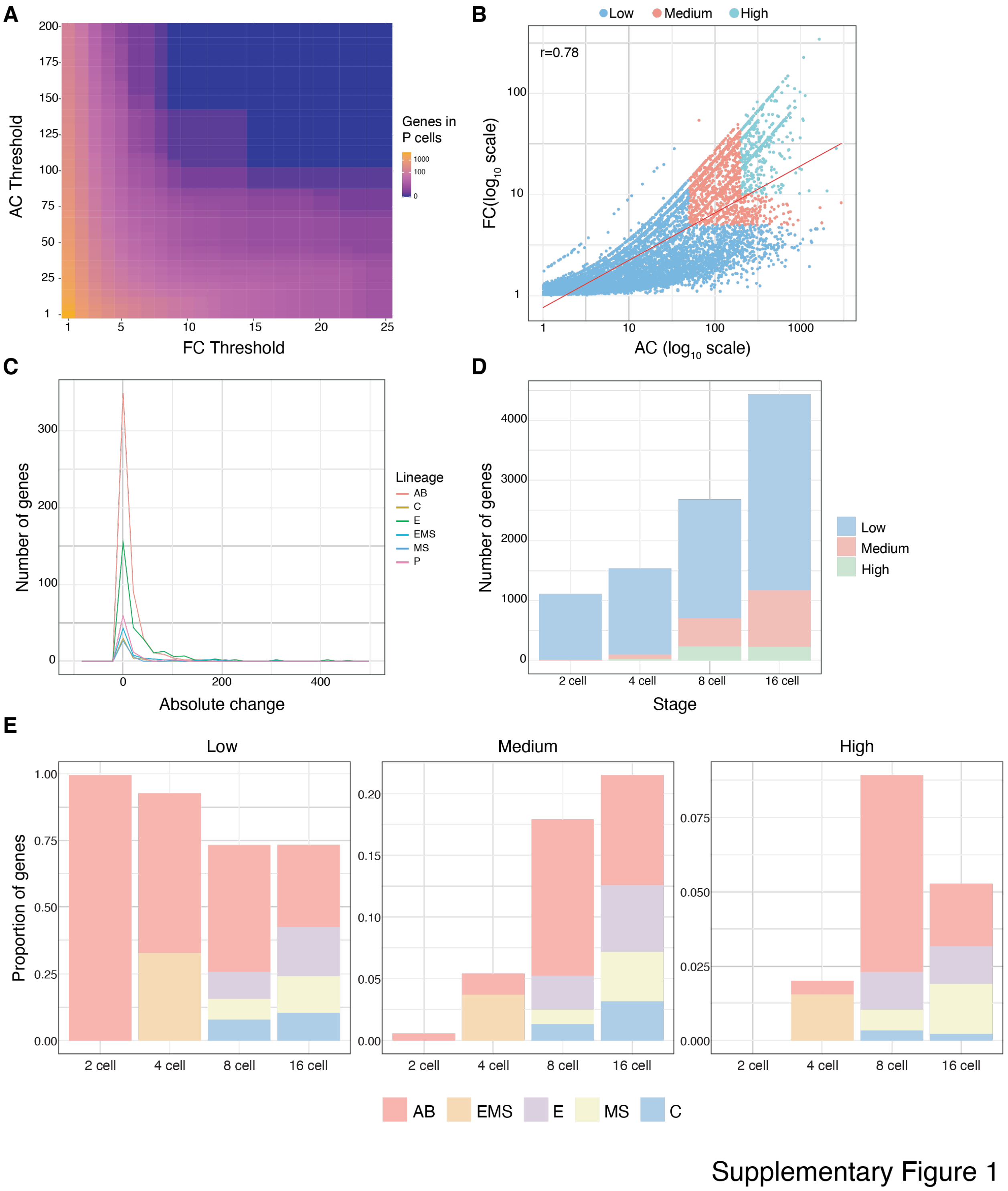


**Supplementary Figure 1 - Related to Fig. 1: Use of absolute change and fold change to categorize accumulation rates of genes.**

**A.** Heatmap of all absolute change (AC) and fold change (FC) as related to the number of genes at each combination of AC and FC in the germline (P cells). **B.** Correlation between AC and FC across different rate categories. **C.** Histogram distribution of positive AC (>1) in different founder lineages. **D.** Number of genes in each rate category at each embryonic cell stage (related to Fig. 1C, D). **E.** Proportion of genes in each rate category at different embryo stages by lineage.

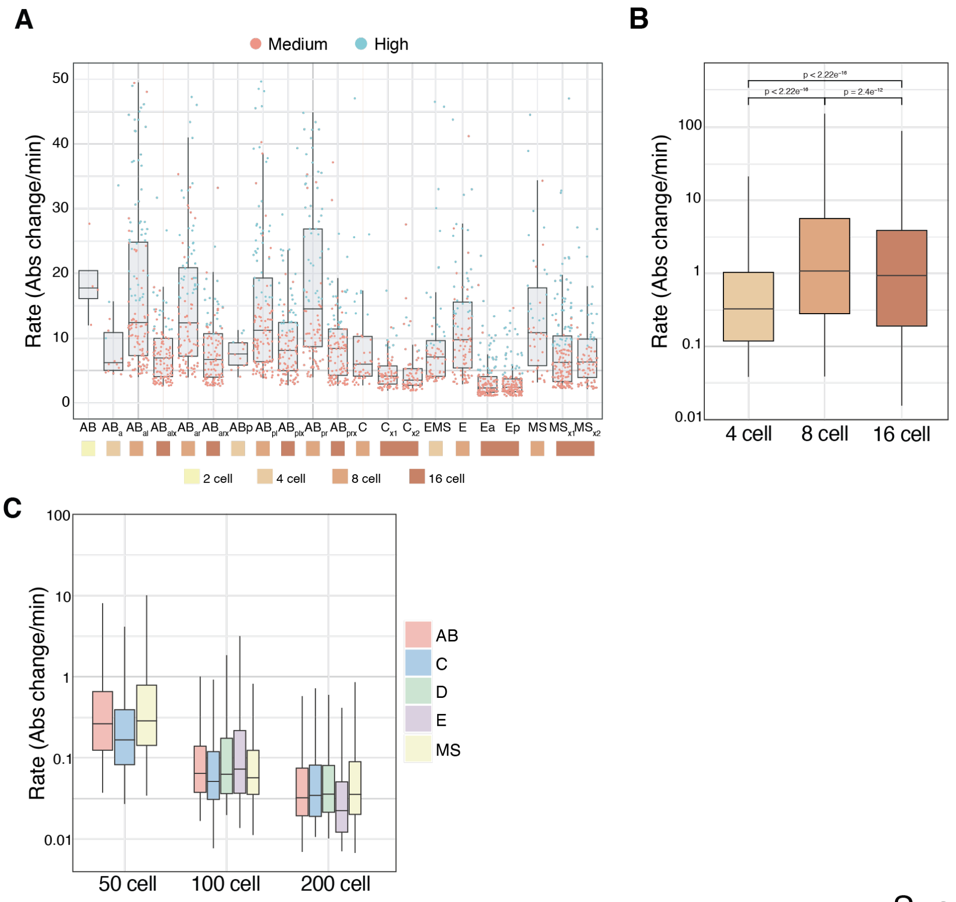


Supplementary Figure 2

**Supplementary Figure 2 - Related to Fig. 2: Rates estimated from scRNA-seq across cell types and lineages.**

**A,B.** Accumulation rate breakdown for high and medium categories by cell type and embryo stage. **C.** Rate breakdown by lineage from the Packer et al. (*33*) data for later embryo stages.

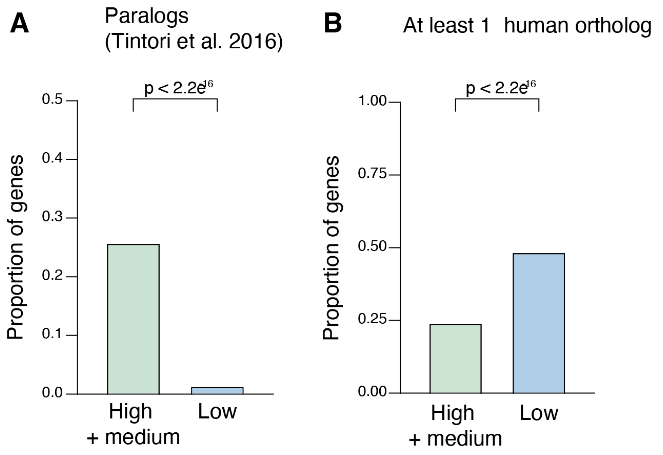


Supplementary Figure 3

**Supplementary Figure 3 – Related to Fig. 3: Paralogs or orthologs in low vs. high-rate genes.**

**A**. Paralogs (Blast result from a e-value threshold of $10^{-15}$) that are also syn-expressed from Tintori et al. (*18*) **B.** Proportion of all high and low-rate genes with at least one human ortholog using Ortholist 2 (*38*).

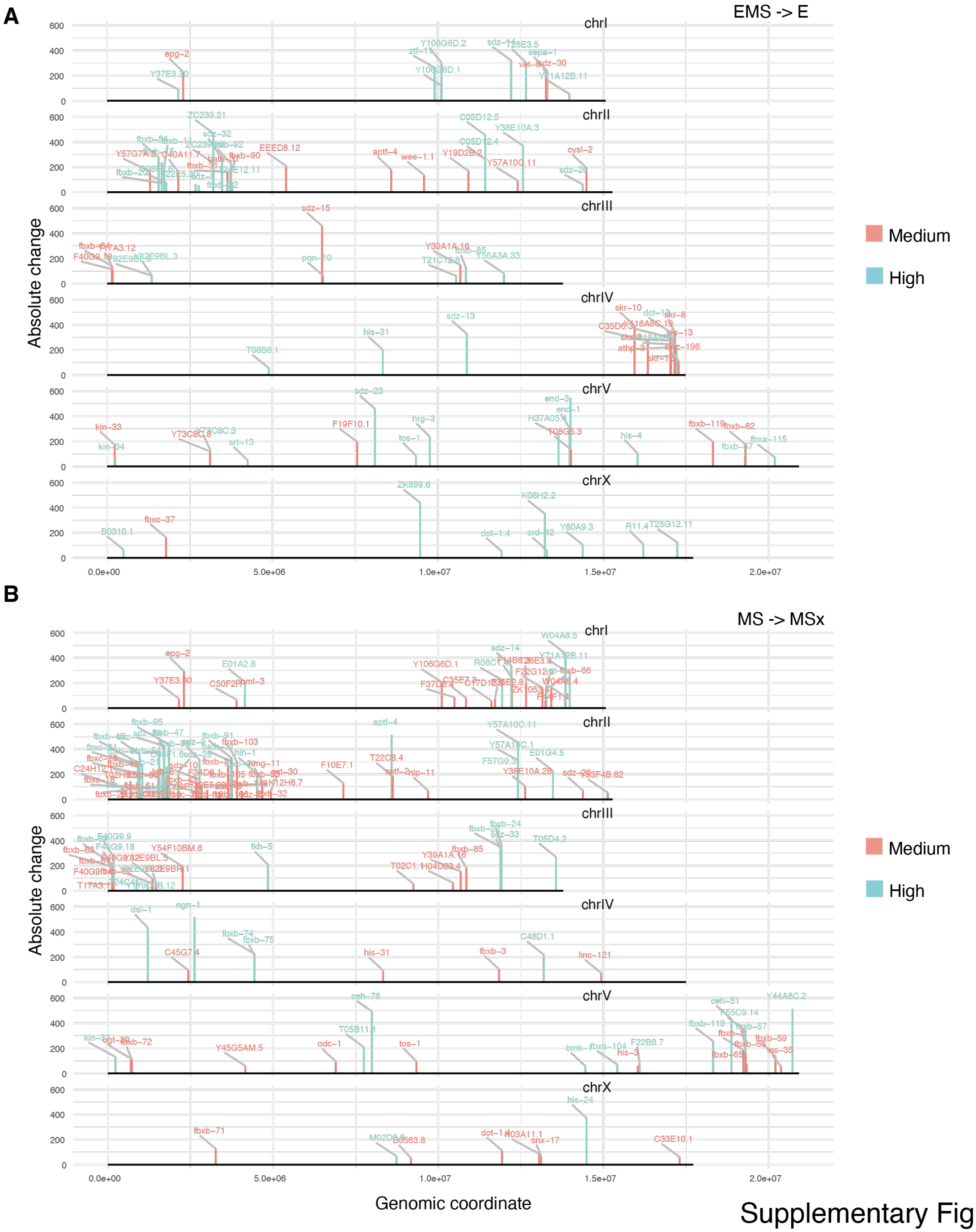


Supplementary Figure 4

**Supplementary Figure 4 – Related to Fig. 4G: Karyogram of clustered genes on all chromosomes for indicated cell types**.

**A.** Karyogram showing absolute change of genes over genomic position (coordinate) for genes in E cells with Fold Change > 5 over the parent EMS cell. **B and C.** Same as A but for MSx1 cell compared to MS parent (B) and for ABprx compared to ABpr parent. Specificity score indicates how specific the expression is to the particular cell (purple is specific to that cell, red is broadly expressed in all cells).

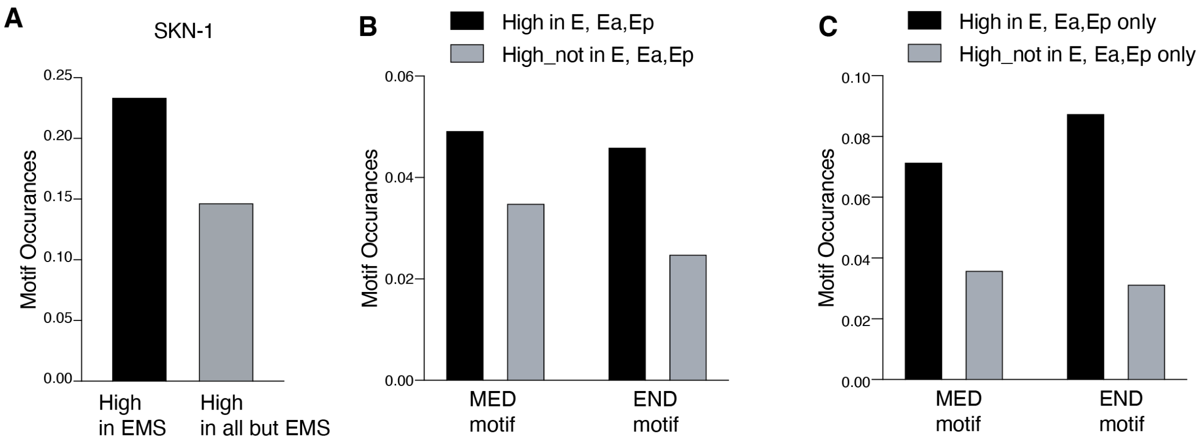


Supplementary Figure 5

**Supplementary Figure 5 – Related to Fig. 5. Motif enrichment for SKN-1, MED and END in the E and MS lineages.**

**A.** SKN-1 motif enrichment in genes expressed in the EMS compared to those expressed in all cells except EMS. **B, C.** END and MED motif enrichment in the E lineage (E, Ea, Ep cells) compared with genes expressed in all but the E lineage. C shows enrichment in genes unique to E *vs.* not.

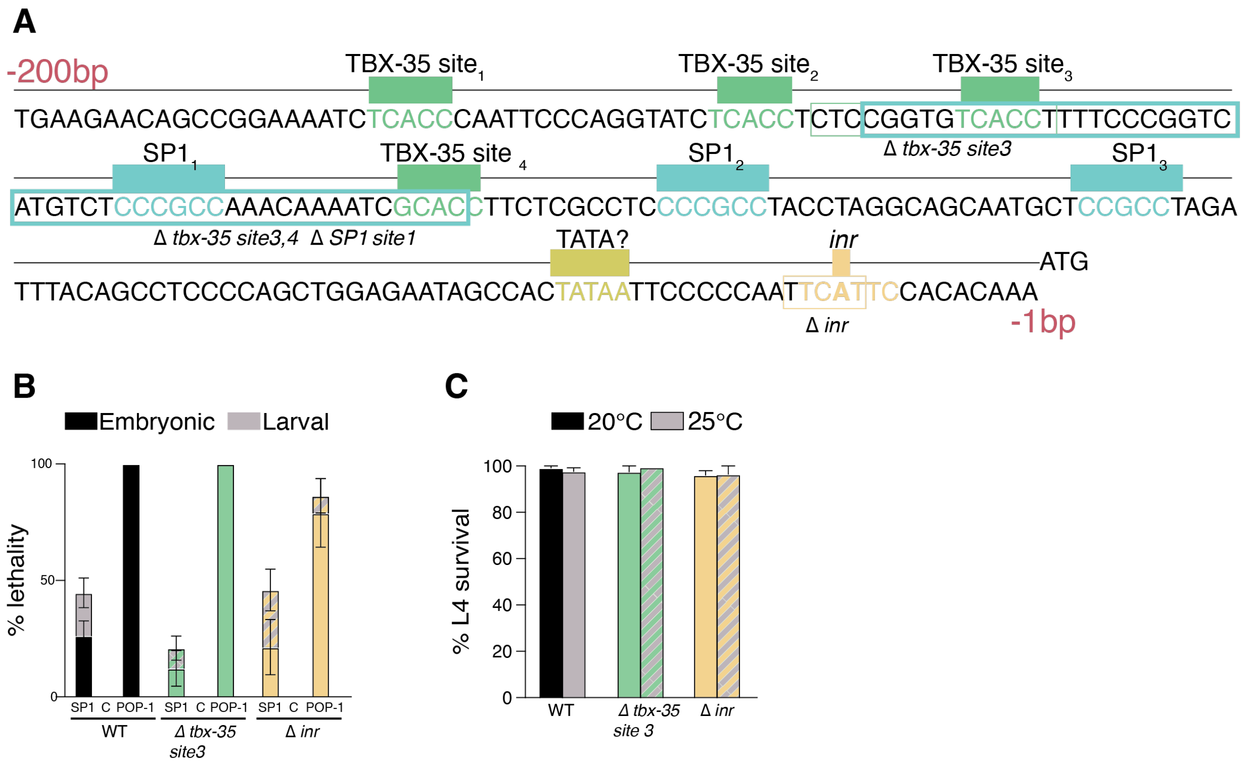


Supplementary Figure 6

**Supplementary Figure 6 – Related to Fig. 6. *ceh-51* promoter region and motif deletion mutants survival**

**A.** Detailed sequence of up to -200bp from the ATG start codon of the *ceh-51* promoter. Motifs indicated in filled colored boxes. Colored boxes around the sequence shows the deleted regions in indicated mutants. **B.** Embryonic and larval lethality after treatment with sptf-3 RNAi (SP1) or pop-1 RNAi as a control, n=at least 4 biological replicates. **C.** Survival in motif mutants at 20°C and 25°C, n=at least 4 biological replicates.
